# Accelerated rTMS for Enhancing Intact Cognition: An Examination of Dose Effects on Electrocortical Indicators of Attention and Working Memory

**DOI:** 10.1101/2025.09.18.676960

**Authors:** Christopher T. Sege, Colin Bowyer, Kevin Caulfield, Donna Roberts, Lisa M. McTeague

## Abstract

**BACKGROUND AND AIMS:** Improvements in cognition (e.g., attention, working memory) are common after repetitive transcranial magnetic stimulation (rTMS) treatment and have also been observed in non-clinical samples. This study investigated whether rTMS can enhance cognitive resilience in individuals who perform in high-stress environments using cognitive measures and electroencephalography (EEG) to explore potential neural mechanisms of rTMS-induced cognitive change.

**METHODS:** 40 college-educated adults not reporting cognitive or psychiatric concerns underwent a 5-day accelerated (10 sessions/ day) rTMS treatment and pre-post cognitive assessment. Participants were assigned to 1 of 10 doses defined as number of active vs. sham stimulation sessions/ day (total active pulses = 3,000 – 30,000). To assess cognitive effects, standard batteries (NIH Toolbox, Spaceflight Cognitive Assessment Tool for Windows [WinSCAT]) were administered – and a subset (n=21) also did an N-Back working memory task with EEG measurement – before, immediately after, and 1 month after rTMS. For all indices, linear and quadratic correlations of pre-to-post-rTMS change with dose were examined to test if an optimal dose was achieved.

**RESULTS:** From pre- to post-rTMS, participants improved in fluid cognition (working memory, processing speed) as measured by NIH toolbox, *t*(39)=8.4, *p*<.001, *d*=1.3, and WinSCAT, *t*(39)=4.1, *p*<.001, *d*=.65; and, improvement in the latter related linearly to rTMS dose, *r*(39)=.40, *p*=.01. Regarding EEG, subjects who received high (6+ active sessions/day) also showed increased amplitudes of an event-related marker of stimulus-directed attention (P300) whether it was elicited by simple task targets, *t*(10)=3.1, *p*=.01, *d*=.95, or task-unrelated noise stimuli played during the task as probes of peripheral attention, *t*(10)=3.6, *p*=.005, *d*=1.1. Additionally, there were linear relationships between change magnitude and rTMS dose for simple task target-related, *r*(20)=.45, *p*=.04, and peripheral noise-related, *r*(20)=.54, *p*=.009, P300s.

**CONCLUSIONS:** rTMS improved fluid reasoning abilities and also changed how dynamic attention is deployed during high-demand challenges as a potential mediator of fluid cognition improvements. While linear dose-response relationships support that changes were rTMS-elicited, absence of a response curve asymptote also suggests that still-higher doses could be warranted to achieve maximal effects in non-clinical samples.

## INTRODUCTION

Repetitive transcranial magnetic stimulation (rTMS), the leading form of non-invasive neuromodulation, has been used as an FDA-approved treatment for refractory major depressive disorder since 2008 (Cohen et al., 2022). rTMS modulates activity in the brain’s cortical surface by rapidly delivering pulses of magnetic field energy to a targeted area under the electro-magnetic coil. In depression treatment, the standard protocol targets dorsolateral prefrontal cortex (dlPFC) – a brain area that is implicated in the pathophysiology of depression (e.g., O’Reardon et al., 2007), can play a key role in the regulation of emotion (e.g., Patel et al., 2023), and is also a central node in a multiple demand network that critically mediates complex cognitive function (McTeague et al., 2016). Related to the latter, improvements in cognitive function are also a commonly reported ancillary benefit of depression treatment (e.g., Martins et al., 2017), which has led to increased research testing the benefits of dlPFC rTMS to cognition in other conditions including neurocognitive conditions like Alzheimer’s Disease (Canter et al., 2016; Weiler et al., 2019; Wu et al., 2022). Even beyond efforts to improve cognition in conditions where it is impaired, the cognitive enhancing effects of rTMS could also be of utility to individuals who have intact cognition but are also susceptible to cognition-impairing effects of performing in extreme settings – for example astronauts who often experience cognitive impairment related to the effects of operating in deep space (Yin et al., 2023). Given this potential use case, then, the current study sought to test the potential for dlPFC rTMS to produce cognitive enhancing effects even in high-performing individuals.

On the strength of its remarkable effectiveness, since its FDA approval rTMS has seen rapid further developments to improve its delivery – including development of patterned pulse delivery patterns that improve efficiency by more closely matching natural brain rhythms (Zrenner & Ziemann, 2024) and accelerated delivery schedules that achieve robust effects in a shorter time than the one session/ day for 5 – 8 weeks standard treatment course (Sonmez et al., 2019; Cole et al., 2020; Caulfield et al., 2022). Development of these protocols not only improves outcomes and efficiency, then, but also raises a need to determine optimal dose-response patterns for rTMS as it is extended beyond depression (e.g., Lewis et al., 2024). Especially in testing rTMS as a tool to enhance cognitive performance in healthy adults, it is critical to determine if an optimal dose can be discovered within the range of what has been tested in accelerated rTMS protocols – or if higher doses are needed in non-clinical populations. Given this need, the primary aim of this study was therefore to examine a dose-response curve of rTMS effects on cognitive performance, as measured with standard neurocognitive testing batteries, in healthy individuals.

In addition, understanding cognitive effects of rTMS in healthy – and indeed also clinical – populations can be further strengthened by testing for neural changes that coincide with observed performance changes and represent potential mediating processes. In addressing this as a second aim, the present study also had a subset of participants do a lab-based cognitive challenge task while brain electroencephalography (EEG) was measured to derive event-related potential (ERP) markers of stimulus-directed attention and task-related cognitive processing. Leveraging EEG sensitivity to temporally granular cognitive/ attentional processing streams (Kappenmann & Luck, 2016), a particular focus here was on a stimulus-elicited P300 ERP component that has been widely studied as an index of how attention is captured by task-relevant or inherently salient stimuli (Donchin, 1980; Polich, 2007). To understand potential TMS impact on how attentional processes are modulated by cognitive demand, then, P300s were elicited in a standard N-Back neurocognitive task (see Owen et al., 2005) where subjects detect matches between stimuli presented in rapid sequence across different levels of working memory load. In addition to measuring task stimulus-elicited P300s, we also measured P300s elicited by attention-grabbing peripheral stimuli presented throughout the task – and as such, used P300 as an index of how both focused and peripheral attention are modulated by cognitive demand, and how rTMS might influence such modulation. By doing this, the second aim of this study was then to determine if we could also detect change in underlying cognitive/ attentional processing brought on by rTMS, and if such change also would demonstrate dose-response effects.

## METHOD

### Participants

Participants in this IRB-approved study were recruited from the population of college-educated adults living around a large Southeastern US Medical Center via flyers, online advertisement, and word-of-mouth referral. Individuals were eligible to if they were between the ages of 18 – 65, had achieved a college degree or higher, and were free of any psychiatric or neurologic (including neurocognitive) disorder. Screening was conducted by study staff prior to enrollment in the study. Study procedures were approved by the local Institutional Review Board and all participants provided written informed consent prior to participation.

### Study Overview

Participation centered around a course of accelerated repetitive rTMS treatment at a randomly assigned dose of active rTMS sessions/ day (see below). Before rTMS, immediately after rTMS, and 1 month after rTMS participants also completed standardized assessments of neurocognitive abilities and neurophysiological correlates. At a baseline visit, participants did consent and then diagnostic interviewing, neurocognitive assessment with standard batteries, and (for participants who elected to do so) neurophysiologic assessment with a working memory task plus electroencephalography (EEG). After baseline, participants came back the next week to undergo rTMS treatment comprising 5 consecutive days of rTMS, and this was immediately followed by repetition of all neurocognitive assessments on the last rTMS day. Finally, participants returned 1 month latter to again repeat all neurocognitive and EEG measures; and at this session, they were finally debriefed and paid after completion of study activities.

### Repetitive Transcranial Magnetic Stimulation (rTMS) Treatment

rTMS treatment comprised an accelerated course of 10 rTMS sessions/ day over 5 days (Fitzgerald et al., 2020). rTMS was delivered using a Magventure MagPro R30 system with a liquid-cooled B65 active/ placebo (A/P) figure-of-eight coil (Farum, Denmark). For all sessions, stimulation intensity was done at 120% resting motor threshold (rMT) – defined as the intensity needed over motor cortex to elicit 5 out of 10 visual twitches in the contralateral anterior pollicis brevis (APB) and determined at baseline for each participant. For treatment, rTMS was targeted to left dlPFC using a standard Beam F3 approach (Beam et al., 2009) and the coil at 45° relative to the sagittal plane. To standardize repeated administration, location and angle at the first session were saved in neuronavigation software (Brainsight Inc., Montreal, Canada) and then replicated at each subsequent treatment day (Caulfield et al., 2022).

In each rTMS session, stimulation was conducted in a standard 50Hz intermittent theta burst (iTBS) pattern of TMS burst trains (3 pulses/ burst; 10 bursts/ train) repeated for 190s (2s on, 7.5s off; 20 trains/ 600 pulses per session). For all subjects, each treatment day comprised 10 consecutive rTMS sessions with 10-min intersession intervals (Fitzgerald et al., 2020). Each participant’s rTMS dose, randomly assigned before treatment, was defined as the number of sessions/ day with *active* stimulation done by conducting rTMS with the B65 A/P coil placed on its active side on the head (doses ranged from 1 – 10 active sessions/ day, i.e., total pulses received over treatment 3,000 – 30,000). Any inactive session involved sham stimulation where the A/P coil was placed on its inactive side and, while stimulation was conducted, surface electrical stimulation was delivered to the scalp at the same rate using the MagPro integrated, synchronized electrical stimulation system and surface electrodes placed under the coil. To minimize psychophysical differences between active and sham sessions, electrical stimulation to the scalp was also done during active sessions. Finally, in addition to blinding participants to condition, TMS operators were also blinded by having a staff member not involved in study visits create each participant order and then orient the coil to the side to be used for the first session so that operators were not aware whether the active or inactive side was used for each session.

### Neurocognitive Assessment

#### Psychiatric Screening – Mini International Neuropsychiatric Interview

The Mini International Neuropsychiatric Interview (M.I.N.I.) for DSM-5 (Sheehan et al., 1998) was conducted to screen for any psychiatric disorder including psychosis or alcohol/ substance use. The M.I.N.I. is a structured interview with separate modules for each DSM-5 psychiatric disorder that include standardized items querying each DSM-5 criterion symptom for that disorder. The M.I.N.I. is exceptionally widely used in psychiatric research and was administered here by a doctoral member of the study team or trained study staff. Unlike all other assessments, the M.I.N.I. was only performed at baseline in this study.

#### Neurocognitive Assessment - NIH Toolbox and Windows Spaceflight Cognitive Assessment Tool

The NIH Toolbox Cognition Battery (Weintraub et al., 2014) is a widely used and standardized brief cognitive assessment battery designed to assess cognitive abilities across a wide range of ability levels. The NIH Toolbox comprises 7 subscales that assess “crystallized” and “fluid” composite dimensions – with the former comprising Picture Vocabulary and Oral Reading subscales that reflect acquired knowledge and the latter comprising Dimensional Card Sort, Flanker Inhibitory Control and Attention, Picture Sequence Memory, List Sorting Working Memory, and Pattern Comparison subscales that reflect theoretically non-acquired cognitive skills. To support its use as a clinical assessment tool, the NIH Toolbox has been normed for the US Population such that composite crystallized and fluid cognition scores can be converted to standardized T scores where 50 represents the US population average. In this study, analyses of rTMS effects focus on standardized crystallized and fluid cognition scores, rather than subscale scores, for clarity.

Complementing the NIH Toolbox (which is designed to assess cognition across a wide array of ability levels and commonly used to assess for neurocognitive disorders), a Windows Spaceflight Cognitive Assessment Tool (WinsCAT; Kane et al., 2005) was also administered here because it was developed to assess cognition specifically in high-performing individuals (Moore et al., 2017). The WinsCAT comprises 5 subscales focused on specifically on fluid intelligence-type abilities such as computation, sustained attentional engagement, task switching, and spatial working memory. In addition to individual subscale scores, WinsCAT subscale scores can be summed to derive a composite score as a general index of cognitive efficiency – and in this study, analyses of rTMS effects focused on this composite for clarity.

### Electroencephalography Recording Session

#### Task-Based Cognitive Assessment – N-Back Attention and Working Memory Task

To test for potential change in neurophysiological correlates of cognition after rTMS, electroencephalography (EEG) was recorded during a standard N-Back task (Gevins & Cutillo, 1993). In the N-back task, participants are shown stimulus sequences (here, shape stimuli presented for 200ms with a 1000 – 2000ms inter-stimulus interval over a 60-s trial) in 3 different target conditions – a 0-back condition, a 1-back condition, and a 2-back condition. Across conditions, the only difference is what defines a “target” during the stimulus sequence – such that in a 0-back condition, a target is simply one type of stimulus (in this study, an “X” shape), in a 1-back condition a target is any stimulus that matches the one presented the turn before, and in a 2-back condition a target is any stimulus that matches one presented *two* turns before (ignoring the intervening stimulus). Before each 60-s trial, the participant is shown a screen indicating the condition of the next trial with the words “0-back,” “1-Back,” or “2-back” (presented for 2s). In all conditions, the participant’s task is then to press one button when a target occurs or another when a non-target occurs, before the next stimulus is shown. In this study, N-Back conditions were presented in counterbalanced pseudorandom order (with each condition occurring no more than twice in a row) and a total of 3 trials per condition. ^1^ On each trial, 20 non-target and 10 target trials were presented such that a total of 60 non-target and 30 targets were presented in total for each condition.

Throughout the N-Back, participants sat 90cm from a computer monitor and task stimuli were presented centrally at 12.5cm x 13.5cm (4 x 4 20’ visual angle from fixation). Along with central stimuli, task-irrelevant noise burst stimuli were also presented throughout as probes of peripheral attention and reflex priming. Noise stimuli were 50ms white noise “bursts” presented with near-instantaneous rise binaurally through in-ear headphones. Noises were presented intermittently in each condition during inter-trial periods, with focus in this presentation on the EEG response elicited by these stimuli as an indicator of peripheral attention.

#### EEG Data Collection

To index event-related potential (ERP) markers of attention and cognitive processing in the N-Back, electroencephalography (EEG) was recorded with a Brain Products© actiCHamp® 32 active-channel system. ActiCap© cloth caps held sensors in 10–20 standard positions on the head. Data were sampled at 500Hz and referenced to a common mode and then sensor Fz.

Impedances were kept below 15 kOhms, and data were not filtered online except for a 250Hz anti-aliasing filter. Along with EEG, electrooculogram data (also sampled at 500Hz) were collected with Ag/ AgCl sensors above and below the right eye and laterally to each eye to correct for gross eye movements during the task.

#### Offline (Post-Task) Data Processing

After collection, EEG processing followed published recommendations (Keil et al., 2014) and for extracting task stimulus and peripheral noise-related event-related potentials (ERPs). Processing was done with Brain Vision Analyzer® and was the same for all N-back shape stimuli and peripheral white noise bursts. Processing began with re-referencing to the average of all sensors and filtering with Butterworth 1/3-amplitude .01Hz (Order 4) – 70Hz (Order 8) cut-off filters and a 60Hz notch filter. Next, EEG was segmented from 200ms before onset – 1000ms after onset of each stimulus and eye movement artifacts were corrected by inputting EEG + electrooculogram data to a blink/ saccade detection algorithm (Gratton et al., 1983).

After eye movement correction, other artifacts were removed based on criteria of overly large (>=200 µV) or small (<=.50 µV) voltages in 200ms windows moving through each segment in 50-ms increments (Keil et al., 2016). Artifact correction was used to identify bad channels that were then interpolated using spherical splines (Junghöfer et al., 2000) unless >2 bad channels were contiguous or >20% of channels were bad on a trial (in which case the trial was removed; Thigpen et al., 2017). After correction, each EEG segment was baseline (100ms – 0ms) corrected and averaged across all trials within each stimulus type.

Following processing as described above, the ERP component of primary interest for both task-relevant and peripheral stimuli – the P300 – was extracted by averaging signal within a sensor and time window where P300 amplitudes were observed to be maximal. Prior studies indicate that canonical P300 positioning and timing varies based on stimulus type such that it is typically centroparietal for visual task stimuli (e.g., Carrasco et al., 2025) and more frontal for auditory startle probes (e.g., Cuthbert et al., 1998). As such, sensor/ time windows used to extract the P300 were allowed to differ for N-Back stimuli versus white noise probes. Actual sensor/ time windows observed for N-Back stimulus and probe P300s are presented in Results.

### Data Analysis

Primary analyses examined pre-to-post-rTMS changes in neurocognitive performance and N-Back performance (i.e., accuracy)/ EEG variables. Initial analyses focused on mixed-effects ANOVAs with a multivariate repeated-measures factor of visit (pre-rTMS, post-rTMS, 1-month follow-up), a between-subjects factor of rTMS dose (defined as high – 6 or more active sessions/ day – or low – 5 or less active sessions/ day), and their interaction. Follow-up paired samples *t*-tests were then used to decompose any significant omnibus effect; and in the event of effects involving visit, linear regression was also used to compute residualized change scores (a post-rTMS – pre-rTMS score and a 1-month follow-up – pre-rTMS score) to be submitted to further analyses of dose effects. In these analyses, complementary strategies were used such that, in one strategy, residualized change scores were compared across high vs. low dose groups with independent *t*-tests; and in another strategy, residualized change scores were also regressed onto continuous rTMS dose to test for linear or quadratic dose-response curves.

Analyses described above were conducted in the same manner for neurocognitive and N-Back variables – and for the latter, analyses were conducted separately for each N-Back condition. Also for N-Back variables, condition effects were tested at baseline to confirm that task performance and ERPs to task variables were modulated across conditions as expected. For baseline analyses of N-Back modulations, task condition (0-back, 1-back, 2-back) was treated as a repeated-measures factor in accordance with published recommendations (Jennings, 1987; Kesselman, 1998). Finally, across all analyses Greenhouse Geisser-corrected degrees of freedom are presented when a sphericity assumption was violated, and effect sizes (partial eta-squared for *f*-statistics, Cohen’s *d* for *t*-statistics) are presented for all tests.

## RESULTS

### Sample

A final sample for this study was 40 participants with mean age 28.1 years (SD=6.7), a roughly equal number of biological women (n=22) and men (n=18), and predominantly reporting non-Hispanic White race/ ethnicity. Diagnostic interview with the MINI Psychiatric Interview confirmed that no participant met diagnostic criteria for a mental health/ substance use disorder.

Of the 40 individuals who completed neurocognitive assessment portions of the study, 26 also completed pre- and post-rTMS EEG assessments while 14 elected not to complete this component. Of the 26 who underwent EEG assessments, 5 (19%) then had their data removed due to poor data/ excessive artifact or technical problems at one or more sessions. Thus, a total of 21 individuals also provided EEG data for analysis in this study.^2^

### Neurocognitive Assessment

#### NIH Toolbox

As expected in this well-educated sample, scores on the NIH Toolbox were above the national average (i.e., average fully corrected T score = 50) for fluid (M=60.3, SD=9.7), *z*(39)= 6.5, *p*<.001, *d*=1.0, and crystallized (M=54.0, SD=8.8), *z*(39)=2.5, *p*=.01, *d*=.41, composite scores and for the total score (M=58.5, SD=8.8), *z*(39)=5.4, *p*<.001, *d*=.85. Next, tests of visit and rTMS dose revealed an effect of visit, *F*(2,76)=57.2, *p*<.001, *η*_*p*_^*2*^*=*.60, but not of dose, *F*(1, 38)=0.9, *p*=.35, *η*_*p*_^*2*^*=*.02, or the Visit X Dose interaction, *F*(2,76)=1.2, *p*=.32, *η*_*p*_^*2*^*=*.01 – and as such, across all participants fluid cognition increased from pre- to post-TMS, *t*(39)=8.4, *p*<.001, *d*=1.3, and was maintained (did not change) to 1-month follow-up, *t*(39)=1.5, *p*=.15, *d*=.23.

Further supporting the lack of a dose effect, follow-up exploration of dose confirmed that: 1) low, *t*(19)=6.7, *p*<.001, *d*=1.5, and high, *t*(19)=5.1, *p*<.001, *d*=1.2, dose groups each showed an increase from pre- to post-rTMS and no further change to 1-month follow-up; 2) residual change scores did not differ across groups for pre-to-post-TMS, *t*(38)=-0.6, *p*=.52, *d*=.20, or post-TMS-to-1-month follow-up, *t*(39)=0.4, *p*=.72, *d*=.12, and; 3) continuous rTMS dose did not correlate with pre-to-post-TMS (linear β=-0.4, *t*[39]=1.3, *p*=.22, *r*=-.20; quadratic β=-0.2, *t*[39]=1.5, *p*=.16) or post-TMS-to-1-month follow-up (linear β=-0.1, *t*[39]=0.3, *p*=.74, *r*=-.06; quadratic β=-0.0, *t*[39]=0.1, *p*=.91) scores. Finally, for crystallized cognition no effect of dose, *F*(1,38)=0.1, *p*=.72, *η*_*p*_^*2*^*=*.00, *or* visit, *F*(2,76)=0.5, *p*=.63, *η*_*p*_^*2*^*=*.00, or of their interaction, *F*(2,76)=0.7, *p*=.49, *η*_*p*_^*2*^*=*.00, arose – and as such, crystallized composite scores did not change from pre- to post-TMS, *t*(39)=0.7, *p*=.48, *d*=.11, or from post-TMS to 1-month follow-up, *t*(39)=0.2, *p*=.82, *d*=.04.

#### WinSCAT

On a total composite score for the WinSCAT, the sample as a whole showed significant change over visits, *F*(2,76)=14.3, *p*<.001, *η*_*p*_^*2*^*=*.27, with no effect of dose, *F*(1,38)=2.8, *p*=.11, *η*_*p*_^*2*^*=*.07, or its interaction with visit, *F*(2,76)=1.9, *p*=.16, *η*_*p*_^*2*^*=*.05. Follow-up analysis of the visit effect indicated an increase from pre-(M=384.1, SD=62.7) to post-(M=415.1, SD=65.6) rTMS, *t*(39)=4.1, *p*<.001, *d*=.65, rTMS, and that this was maintained at 1-month follow-up (M=419.4, SD=58.1) such that post-rTMS and follow-up did not differ, *t*(39)=0.5, *p*=.59, *d*=.09 (Fig. 1).

**Figure 1.**
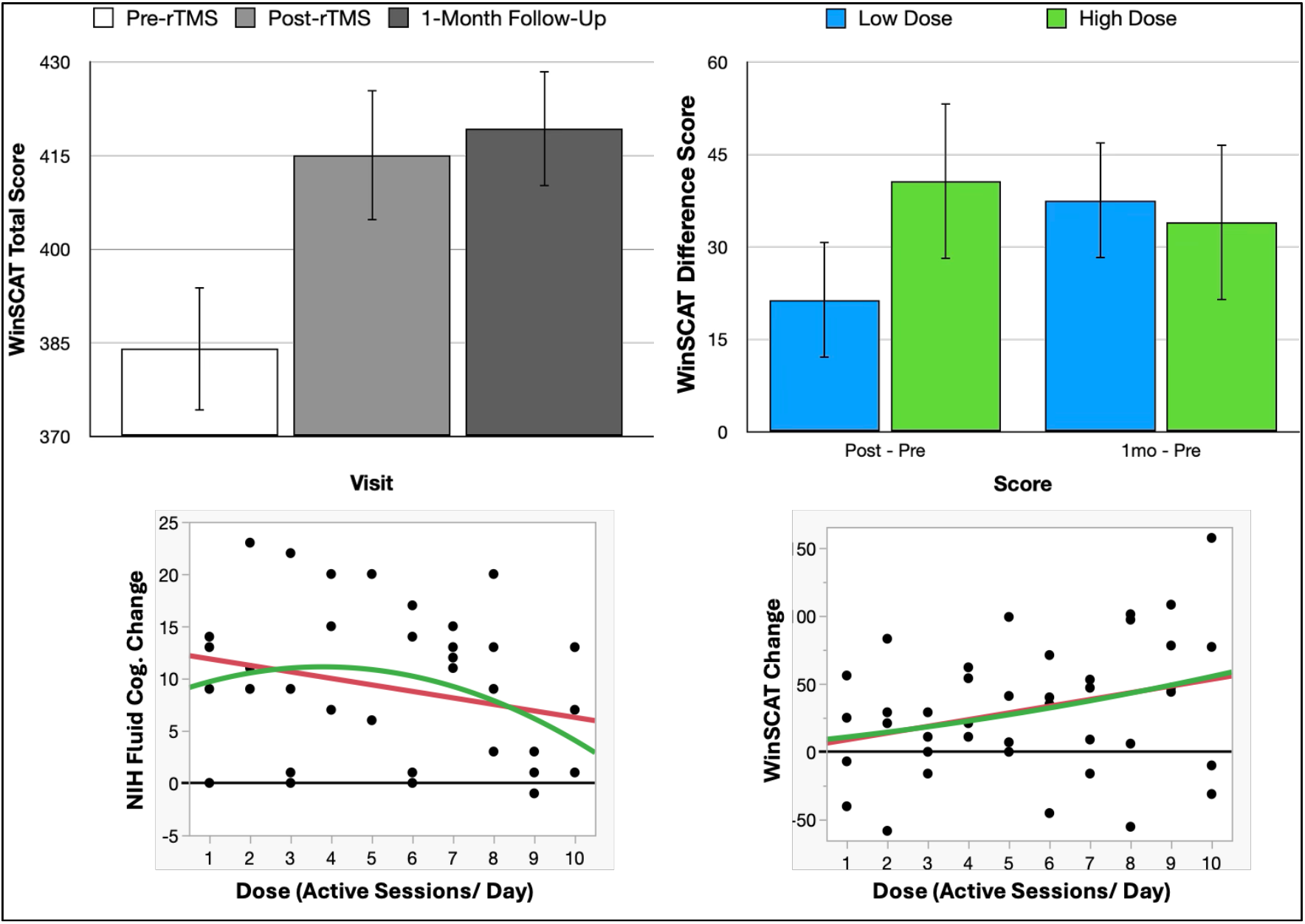
Cognitive changes from before to after rTMS treatment. Top left depicts change in WinSCAT total score across visits, and top right panel depicts WinSCAT change for low and high rTMS dose groups. Bottom panels depict linear (red) and quadratic (green) dose trends for NIH fluid cognition (left) and WinSCAT total (right) post-– pre-rTMS change scores.

Further exploration of potential effects of dose then did provide some evidence for distinct patterns such that: 1) for a low rTMS dose group, WinsCAT score increased from pre- to post-rTMS, *t*(19)=2.5, *p*=.02, *d*=.56, and again from post-rTMS to 1-month follow-up, *t*(19)=2.3, *p*=.03, *d*=.52, whereas; 2) for a high dose rTMS group, a marked increase from pre- to post-rTMS, *t*(19)=3.3, *p*=.004, *d*=.73, was maintained, with no further increase, *t*(19)=0.7, *p*=.49, *d*=.16, to 1-month follow-up (Fig 1). Comparing residualized change scores across groups further revealed a difference in pre-to-post-TMS change that approached significance, *t*(38)=1.8, *p*=.08, *d*=.58, and that groups did not differ in their pre-TMS, *t*(38)=1.3, *p*=.21, *d*=.40, or 1-month follow-up, *t*(38)=0.9, *p*=.51, *d*=.22, scores but did differ significantly in post-TMS scores, *t*(38)= 2.2, *p*=.03, *d*=.71. Contextualizing group comparisons, continuous rTMS dose also correlated linearly with pre-to-post-TMS residualized change (linear β=6.3, *t*[39]=2.7, *p*=.01, *r*=.40; quadratic β=0.2, *t*[39]=0.2, *p*=.87) but not with post-TMS-to-1-month follow-up change (linear β=0.2, *t*[39]=0.1, *p*=.93, *r*=.02; quadratic β=-0.4, *t*[39]=0.6, *p*=.59). Consistent with change score results, dose also correlated linearly with post-rTMS (linear β=10.4, *t*[39]=3.2, *p*=.003, *r*=.46; quadratic β=0.1, *t*[39]=0.1, *p*=.92) but not pre-rTMS (linear β=5.5, *t*[39]=1.6, *p*=.12, *r*=.25; quadratic β=-0.0, *t*[39]=0.0, *p*=.98) or 1-month follow-up (linear β=4.2, *t*[39]=1.3, *p*=.20, *r*=.21; quadratic β=-0.5, *t*[39]=0.4, *p*=.71) scores.

### N-Back Task

#### Performance Accuracy

At baseline, N-Back task accuracy was modulated across conditions, *F*(2,40)=5.1, *p*= .01, *η*_*p*_^*2*^*=*.20 – such that participants were more accurate in a 1-back condition (M= 89.4%, SD=13.7%) than 0-back (M=85.2%, SD=21.0%), *t*(20)=1.9, *p*=.08, *d*=.41, or 2-back (M= 82.8%, SD=13.3%), *t*(20)=4.4, *p*<.001, *d*=1.0, conditions. 1- and 2-back conditions did not differ from each other, *t*(20)=1.0, *p*=.33, *d*=.22. Next, analysis of visit and dose effects indicated marginal effects of visit, *F*(2,38)=3.2, *p*=.08, *η*_*p*_^*2*^*=*.14, and dose, *F*(1,19)=4.1, *p*=.06, *η*_*p*_^*2*^*=*.18, along with a non-significant interaction, *F*(2,38)=1.1, *p*=.33, *η*_*p*_^*2*^*=*.06, for the 0-back condition. Follow-up of a visit effect showed that 0-back hit percentage increased from pre- to post-rTMS, *t*(20)=2.3, *p*= .03, *d*=.51, and there was then no further increase to 1-month follow-up, *t*(20)=0.9, *p*=.37, *d*=.20. Next, following up the dose effect indicated marginal increased accuracy in the high-dose group pre-rTMS, *t*(19)=1.8, *p*=.10, *d*=.77, and 1-month follow-up, *t*(19)=1.8, *p*=.09, *d*=.77, along with no group differences in post-rTMS or 1-month follow-up residual change score. No bivariate correlation of raw scores or residualized change scores with dose were significant (all *p*s > .12).

Next, for a 1-back condition no effect of visit, *F*(2,38)=0.8, *p*=.44, *η*_*p*_^*2*^*=*.04, dose, *F*(1,19)= 2.0, *p*=.17, *η*_*p*_^*2*^*=*.10, or their interaction, *F*(2,38)=1.2, *p*=.31, *η*_*p*_^*2*^*=*.06, arose. For the 2-back condition, then, there was no dose, *F*(1,19)=1.8, *p*=.20, *η*_*p*_^*2*^*=*.09, or Dose X Visit, *F*(2,38)=0.9, *p*=.41, *η*_*p*_^*2*^*=*.01, effect but the effect of visit was significant, *F*(2,38)=4.8, *p*=.01, *η*_*p*_^*2*^*=*.20; and following this up, paired *t*-tests revealed specific improvement from pre-TMS to 1-month follow-up, *t*(20)=3.4, *p*=.003, *d*=.73, whereas performance immediately after rTMS did not differ from before rTMS, *t*(20)=1.4, *p*=.18, *d*=.30, or 1-month follow-up, *t*(20)=1.5, *p*=.15, *d*=.33. Further contextualizing this, comparing residualized change scores across high vs. low rTMS dose groups did not indicate a difference for any score, including a pre-rTMS-to-1-month score, *t*(20)=0.0, *p*=.97, *d*=.01; and, rTMS dose did not correlate with any 2-back score (all *p*s > .16).

#### N-Back Target P300

As expected, N-back target stimuli elicited a robust P300 ERP component that was maximal from 370 – 470ms at centroparietal sites (Cz, CP1, CP2, Pz). Also as expected, target P300 was enhanced relative to non-targets across all conditions (see Fig 2), but the degree of enhancement was modulated by condition, *F*(2,40)=5.1, *p*=.01, *η*_*p*_^*2*^*=* .20, such that P300 (target – non-target) amplitude was larger in the 0-back condition than in the 1-back, *t*(20)=2.0, *p*=.06, *d*=.44, and 2-back, *t*(20)=2.9, *p*=.01, *d*=.62, conditions. P300s did not differ across the 1-back and 2-back conditions, *t*(20)=1.2, *p*=.24, *d*=.26.

**Figure 2.**
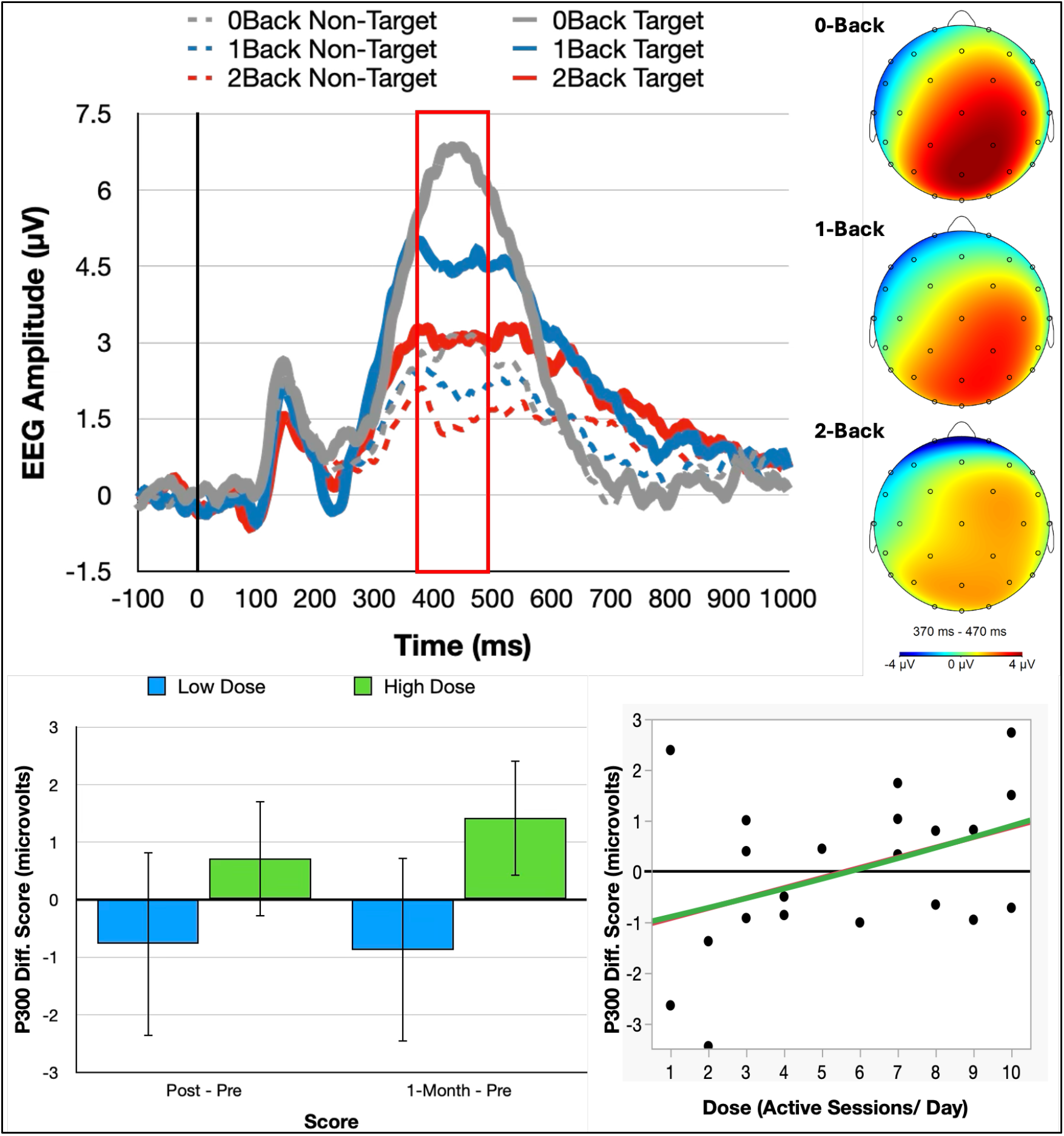
N-Back task stimulus ERP. Top panel depicts ERP waveforms for target and non-targets in each condition (P300 window indicated by red box), and topographic plots of P300 (target – non-target) amplitudes for each condition. Bottom left panel depicts change scores for low and high rTMS dose groups. Bottom right panel depicts linear (red) and quadratic (green) dose trends for P300 1-month follow-up - pre-rTMS change score.

After analysis of baseline modulation patterns, testing change across visits in the 0-back task condition indicated no effect of visit, *F*(2,38)=0.2, *p*=.82, *η*_*p*_^*2*^*=* .01, or dose, *F*(1,19)=0.4, *p*=.56, *η*_*p*_^*2*^*=* .02, but a Visit X Dose interaction that approached significance, *F*(2,38)=2.9, *p*=.08, *η*_*p*_^*2*^*=*.13. Follow-up indicated no change from pre- to post-rTMS, *t*(9)=0.6, *p*=.55, *d*=.20, or post-rTMS to 1-month follow-up, *t*(9)=0.1, *p*=.89, *d*=.05, in the low dose group; but in the high dose group, P300 amplitudes did increase from pre-TMS to 1-month follow-up, *t*(10)=3.1, *p*=.01, *d*=.95. Further contextualizing these differences, comparing residual change scores across high vs. low rTMS dose groups revealed that those in the high group had larger pre-rTMS-to-1-month follow-up change scores than did those in the low group, *t*(19)=2.3, *p*=.04, *d*=1.0; and additionally, rTMS dose correlated linearly with the 1-month follow-up – pre-rTMS change score (linear β=0.4, *t*[20]=2.2, *p*=.04, *r*=.45; quadratic β=0.02, *t*[20]=0.3, *p*=.78).

Unlike the 0-back condition, analysis of the 1-back indicated no effect of visit, *F*(2,38)= 0.8, *p*=.47, *η*_*p*_^*2*^*=* .01, dose, *F*(2,38)=0.5, *p*=.48, *η*_*p*_^*2*^*=* .03, or their interaction, *F*(2,38)= 0.3, *p*=.78, *η*_*p*_^*2*^*=* .01. Similarly, for the 2-back condition there was also no effect of visit, *F*(2,38)=0.9, *p*=.39, *η*_*p*_^*2*^*=* .05, dose, *F*(2,38)= 0.1, *p*=.79, *η*_*p*_^*2*^*=* .004, or the interaction, *F*(2,38)= 0.3, *p*=.67, *η*_*p*_^*2*^*=* .02.

#### N-Back Peripheral Stimulus (Startle Probe) P300

Peripheral white noise bursts presented during the N-back task also elicited a robust P300 ERP component – in this case maximal from 330 – 400ms and at frontocentral sites (Fz, FC1, FC2, Cz; see Fig. 3). Like N-back target P300s, probe P300 amplitudes were modulated across conditions, *F*(2,38)=5.1, *p*=.01, *η*_*p*_^*2*^*=* .20 – in this case so that P300s were enhanced in 1-back, *t*(20)=2.4, *p*=.03, *d*=.52, and 2-back, *t*(20)=3.3, *p*=.004, *d*=.71, conditions relative to 0-Back, and 1-back and 2-back conditions did not differ, *t*(20)=0.1, *p*=.91, *d*=.02.

**Figure 3.**
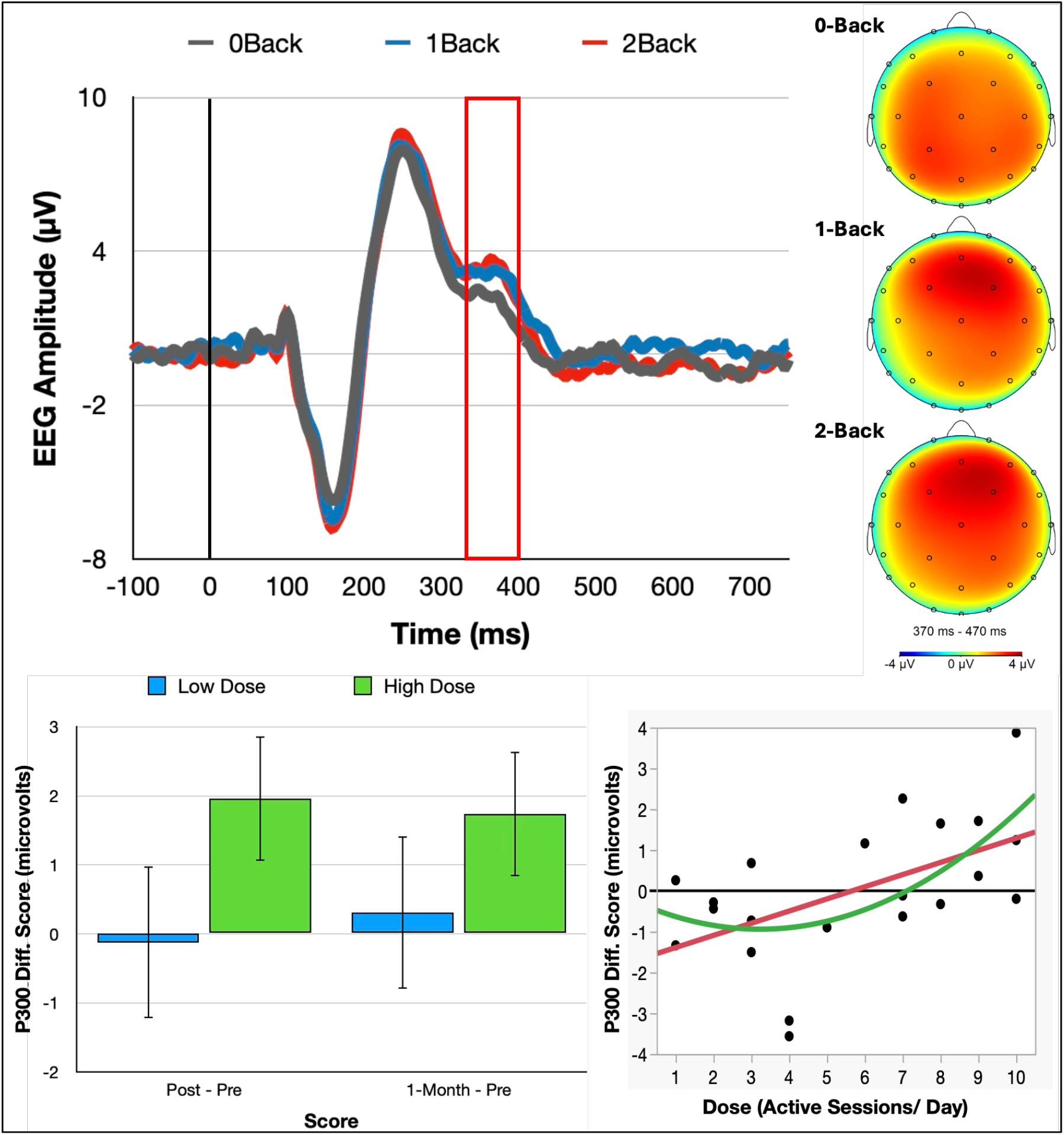
Peripheral stimulus ERP. Top depicts waveforms across conditions (P300 window indicated by red box), and topographic plots of amplitudes for each condition. Bottom left panel depicts change scores for low and high rTMS dose groups. Bottom right panel depicts linear (red) and quadratic (green) dose trends for P300 post-rTMS - pre-rTMS change score.

After analysis of baseline modulation patterns, change testing for each condition indicated significant change across visits for the 0-back condition, *F*(2,38)=5.8, *p*=.01, *η*_*p*_^*2*^*=* .23, that was qualified by an interaction with dose group, *F*(2,38)=5.2, *p*=.02, *η*_*p*_^*2*^*=* .21. Following up this interaction, paired *t*-tests in each group revealed no changes from pre- to post-rTMS, *t*(9)=0.3, *p*=.78, *d*=.09, or from post-rTMS to 1-month follow-up, *t*(9)=0.9, *p*=.39, *d*=.28, in the low-dose group; while in the high group, there was a significant increase in P300 amplitude at post-rTMS, *t*(10)=4.8, *p*<.001, *d*=1.4, and 1-month follow-up, *t*(10)=3.6, *p*=.005, *d*=1.1, relative to pre-rTMS. Contextualizing these different patterns, comparison of residualized change scores across high vs. low rTMS dose groups further revealed that individuals in a high group had larger pre-to-post-rTMS changes in startle P300, *t*(19)=3.5, *p*=.002, *d*=1.5, and approached larger changes from pre-to-1-month follow-up, *t*(19)=1.8, *p*=.09, *d*=.78, compared to those in the low group. In addition, continuous rTMS dose correlated with the post-rTMS – pre-rTMS change score (linear β=0.3, *t*[20]=2.9, *p*=.009, *r*=.54; quadratic β=0.06, *t*[20]=1.4, *p*=.17) and its correlation with 1-month follow-up – pre-rTMS change score approached significance, (linear β=0.25, *t*[20]=1.8, *p*=.08, *r*=.39; quadratic β=0.06, *t*[20]=0.6, *p*=.55) for the 0-back condition.

Analyses of 1-back and 2-back conditions indicated different patterns than for 0-back such that no effect of visit arose for 1-back, *F*(2,38)= 0.5, *p*=.62, *η*_*p*_^*2*^*=* .01, or 2-back, *F*(2,38)= 0.9, *p*=.42, *η*_*p*_^*2*^*=* .04, and there was also no Visit X Dose interaction for 1-back, *F*(2,38)= 1.3, *p*=.29, *η*_*p*_^*2*^*=* .06, or 2-back, *F*(2,38)= 1.4, *p*=.25, *η*_*p*_^*2*^*=* .02, conditions; but at the same time, a main effect of dose approached significance for 1-back, *F*(2,38)= 3.5, *p*=.08, *η*_*p*_^*2*^*=* .16, and was significant for 2-back, *F*(2,38)= 10, *p*=.005, *η*_*p*_^*2*^*=* .35. Follow-up of dose effects in 1- and 2-back conditions revealed a low-dose vs. high-dose group difference for the 1-back condition only at 1-month follow-up, *t*(19)=2.3, *p*=.03, *d*=1.0; while for the 2-back condition, group differences tended toward significance pre-rTMS, *t*(19)=1.8, *p*=.09, *d*=.78, and were significant at post-rTMS, *t*(19)=2.8, *p*=.01, *d*=1.2, and 1-month follow-up, *t*(19)=3.1, *p*=.006, *d*=1.4. Qualifying this, exploration of group differences in residualized change scores also revealed no differences for 1-back but that groups did differ for post-rTMS – pre-rTMS, *t*(19)=1.8, *p*=.09, *d*=.79, and 1-month follow-up – pre-rTMS, *t*(19)=2.2, *p*=.04, *d*=.96, scores in the 2-back condition. In addition, dose also correlated linearly with 2-back post-rTMS – pre-rTMS (linear β=0.3, *t*[20]=2.0, *p*=.05, *r*=.43; quadratic β=-0.01, *t*[20]=0.2, *p*=.83) and 1-month follow-up – pre-rTMS (linear β=0.4, *t*[20]=2.4, *p*=.03, *r*=.49; quadratic β=0.02, *t*[20]=0.3, *p*=.74) change scores; and, not with P300s at pre-rTMS baseline in 0-back, (linear β=-0.1, *t*[20]=0.5, *p*=.63, *r*=-.12; quadratic β=0.03, *t*[20]= 0.6, *p*=.59), 1-back (linear β=0.2, *t*[20]=1.2, *p*=.25, *r*=.27; quadratic β=-0.02, *t*[20]=0.2, *p*=.82), or 2-back (linear β=0.2, *t*[20]=1.4, *p*=.19, *r*=.31; quadratic β=-0.03, *t*[20]=0.5, *p*=.62) conditions.

## DISCUSSION

To advance our understanding of rTMS as a tool to enhance cognition in healthy adults, the present study tested the relationship of cognitive effects to TMS dose and explored possible neurofunctional mechanisms of these effects using brain electrophysiology measures. In addressing its aims, this study leveraged advancements in *accelerated* rTMS to deliver a high number of rTMS sessions per day, with dose operationalized as the number of sessions each day that delivered active (versus psychophysically matched non-active surface) stimulation. Using this approach, results confirmed pre-to-post improvements in fluid cognitive abilities (e.g., processing speed, working memory) that were maintained to 1-month follow-up on NIH Toolbox and WinSCAT measures, and they found a linear dose-response relationship of pre-to-post change for the WinSCAT measure. Adding to this, an electrophysiological index of attention deployment – the P300 event-related potential (Donchin, 1980; Polich, 2007) – showed change from pre-rTMS to 1-month follow-up, and dose-response effects for this change, for P300s elicited by simple target stimuli in an N-Back task (i.e., “X” shape targets in the 0-back condition) or by peripheral (task-irrelevant) white noise bursts. Taken together, then, results further support that rTMS can produce improvements especially in fluid cognitive abilities in healthy adults, and also provide initial evidence that an impact on how stimulus-directed attention is deployed in dynamic demand contexts might be a route by which rTMS produces such improvements.

The elucidation of a dose-response relationship between cognitive effects and number of rTMS sessions (and thus total number of rTMS pulses delivered) extends evidence for cognition-enhancing effects of TMS even in cognitively intact individuals (see Ebrahimzadeh et al., 2025) – such that dose relationships further support that changes are induced by rTMS rather than practice-related. The fact that a dose-response relationship arose specifically for WinSCAT could be due to the fact this measure was designed specifically to be challenging to high-performing individuals like the college-educated individuals recruited for the study (Moore et al., 2017) – and thus is less susceptible than NIH Toolbox to large and consistent practice effects that could obscure rTMS-induced changes. Further contextualizing dose-response effects, the fact that this relationship was linear for the WinSCAT (and task-based electrophysiology indicators) suggests that even stronger effects might be produced by still-higher doses of rTMS. Thus, while this study is the largest bolus of rTMS treatment ever delivered, in published research, to healthy individuals, future research should continue to explore even higher doses to determine where an asymptote in the dose-response curve – indicating optimal dose to produce cognitive effects – is met.

Contextualizing effects on the neurocognitive batteries, electrophysiological findings then provide initial support for further exploring changes in online attention deployment as a potential mediator of cognition-enhancing effects of rTMS. The P300 event-related potential component analyzed here is widely considered, based on a long history of research, to capture preferential neural/ attentional processing of stimulus events that are *salient* because they are task-relevant (e.g., Heinze et al., 1990), novel (e.g., Luck et al., 2009), or inherently detected to be relevant (e.g., Cuthbert et al., 1998). In this study, enhancement of P300s for task-relevant discrete targets *and* peripheral but also inherently salient stimuli could suggest that rTMS broadens the capacity to detect salient stimuli in demanding contexts as a substrate to better performance in response to these stimuli. While more research is needed to confirm this speculation, nonetheless a dose-response relationship also again supports that P300 changes observed here were indeed rTMS-induced – and points to an opportunity to explore even higher doses to potentially produce stronger neurofunctional effects.

To build on findings from this study, further research must address limitations of current work that leave still-open questions. Most critically, future work must replicate and extend current methods in larger samples – which would, for one, allow for direct testing of mediational relationships between neurocognitive and electrophysiological indices as speculated above. In addition, future work must also explore higher doses of rTMS by, if possible, increasing number of pulses delivered within the same (1-week) temporal span – research needed to find an asymptote in the dose-response curve that would indicate the upper limit of potential for enhancement-producing effects. Finally, to understand electrocortical effects future research must also explore additional task contexts beyond the N-Back – with perhaps particular focus on multiple-demand/ divided attention tasks that could further explicate if rTMS indeed does widen directed attention capacity as a mechanism for producing its cognitive enhancing effects.

Even before future work, current research contributes to our understanding of rTMS as a tool for enhancing cognition in high-performing individuals by demonstrating dose-response relationships and beginning to point to neurofunctional mechanisms as candidates for further study. Replicating and extending these results could greatly support development of rTMS as a tool for improving cognition in neurocognitive disorder and enhancing cognitive resilience in healthy individuals. For the latter population, such a tool could be especially useful for those who operate in high-stress environments as a tool for protecting against the cognitive decrements that such environments produce.

## Footnotes

1 A high-vs. low-stress manipulation was also employed with the N-Back task such that trials were presented with a blue or yellow background (luminance-matched) that indicated whether an aversive stimulus (a loud human scream) might also occur during that trial. Because statistical analyses revealed no modulation of task variables or rTMS eBects as a function of the stress manipulation, this manuscript only presents data for low-stress trials that are more consistent with the canonical N-Back task to maintain clarity of presentation.

2 3 participants had pre and post, but not 1-month follow-up, cognitive battery (NIH Toolbox, WinSCAT) data; and, of the 21 individuals with pre and post EEG, 2 did not have EEG data at a 1-month follow-up. To maintain sample size for pre-to-post rTMS comparison, missing data at 1-month follow-up were interpolated from the post-rTMS visit for each index. Patterns of results did not diBer in any case when subjects with interpolated data were removed.

## References Cited

Cohen SL, Bikson M, Badran BW, George MS. A visual and narrative timeline of US FDA milestones for Transcranial Magnetic Stimulation (TMS) devices. Brain Stimul. 2022 Jan-Feb;15(1):73–75. doi: 10.1016/j.brs.2021.11.010. Epub 2021 Nov 11. PMID: 34775141; PMCID: PMC8864803.

O’Reardon JP, Solvason HB, Janicak PG, Sampson S, Isenberg KE, Nahas Z, McDonald WM, Avery D, Fitzgerald PB, Loo C, Demitrack MA, George MS, Sackeim HA. Efficacy and safety of transcranial magnetic stimulation in the acute treatment of major depression: a multisite randomized controlled trial. Biol Psychiatry. 2007 Dec 1;62(11):1208–16. doi: 10.1016/j.biopsych.2007.01.018. Epub 2007 Jun 14. PMID: 17573044.

Patel M, Teferi M, Casalvera A, Lynch K, Nitchie F, Makhoul W, Oathes DJ, Sheline Y, Balderston NL. Interleaved TMS/fMRI shows that threat decreases dlPFC-mediated top-down regulation of emotion processing. medRxiv [Preprint]. 2023 Nov 11:2023.11.11.23298414. doi: 10.1101/2023.11.11.23298414. PMID: 37986856; PMCID: PMC10659468.

McTeague LM, Goodkind MS, Etkin A. Transdiagnostic impairment of cognitive control in mental illness. J Psychiatr Res. 2016 Dec;83:37–46. doi: 10.1016/j.jpsychires.2016.08.001. Epub 2016 Aug 5. PMID: 27552532; PMCID: PMC5107153.

Martins AR, Fregni F, Simis M, Almeida J. Neuromodulation as a cognitive enhancement strategy in healthy older adults: promises and pitfalls. Neuropsychol Dev Cogn B Aging Neuropsychol Cogn. 2017 Mar;24(2):158–185. doi: 10.1080/13825585.2016.1176986. Epub 2016 Apr 28. PMID: 27123674.

Canter RG, Penney J, Tsai LH. The road to restoring neural circuits for the treatment of Alzheimer’s disease. Nature. 2016 Nov 10;539(7628):187–196. doi: 10.1038/nature20412. PMID: 27830780.

Weiler M, Stieger KC, Long JM, Rapp PR. Transcranial Magnetic Stimulation in Alzheimer’s Disease: Are We Ready? eNeuro. 2020 Jan 7;7(1):ENEURO.0235-19.2019. doi: 10.1523/ENEURO.0235-19.2019. PMID: 31848209; PMCID: PMC6948923.

Wu X, Ji GJ, Geng Z, Wang L, Yan Y, Wu Y, Xiao G, Gao L, Wei Q, Zhou S, Wei L, Tian Y, Wang K. Accelerated intermittent theta-burst stimulation broadly ameliorates symptoms and cognition in Alzheimer’s disease: A randomized controlled trial. Brain Stimul. 2022 Jan-Feb;15(1):35–45. doi: 10.1016/j.brs.2021.11.007. Epub 2021 Nov 6. PMID: 34752934.

Yin Y, Liu J, Fan Q, Zhao S, Wu X, Wang J, Liu Y, Li Y, Lu W. Long-term spaceflight composite stress induces depression and cognitive impairment in astronauts-insights from neuroplasticity. Transl Psychiatry. 2023 Nov 8;13(1):342. doi: 10.1038/s41398-023-02638-5. PMID: 37938258; PMCID: PMC10632511.

Zrenner C, Ziemann U. Closed-Loop Brain Stimulation. Biol Psychiatry. 2024 Mar 15;95(6):545–552. doi: 10.1016/j.biopsych.2023.09.014. Epub 2023 Sep 22. PMID: 37743002; PMCID: PMC10881194.

Sonmez AI, Camsari DD, Nandakumar AL, Voort JLV, Kung S, Lewis CP, Croarkin PE. Accelerated TMS for Depression: A systematic review and meta-analysis. Psychiatry Res. 2019 Mar;273:770–781. doi: 10.1016/j.psychres.2018.12.041. Epub 2018 Dec 7. PMID: 31207865; PMCID: PMC6582998.

Cole EJ, Stimpson KH, Bentzley BS, Gulser M, Cherian K, Tischler C, Nejad R, Pankow H, Choi E, Aaron H, Espil FM, Pannu J, Xiao X, Duvio D, Solvason HB, Hawkins J, Guerra A, Jo B, Raj KS, Phillips AL, Barmak F, Bishop JH, Coetzee JP, DeBattista C, Keller J, Schatzberg AF, Sudheimer KD, Williams NR. Stanford Accelerated Intelligent Neuromodulation Therapy for Treatment-Resistant Depression. Am J Psychiatry. 2020 Aug 1;177(8):716–726. doi: 10.1176/appi.ajp.2019.19070720. Epub 2020 Apr 7. PMID: 32252538.

Caulfield KA, Fleischmann HH, George MS, McTeague LM. A transdiagnostic review of safety, efficacy, and parameter space in accelerated transcranial magnetic stimulation. J Psychiatr Res. 2022 Aug;152:384–396. doi: 10.1016/j.jpsychires.2022.06.038. Epub 2022 Jun 28. PMID: 35816982; PMCID: PMC10029148.

Lewis CP, Nakonezny PA, Sonmez AI, Ozger C, Garzon JF, Camsari DD, Yuruk D, Romanowicz M, Shekunov J, Zaccariello MJ, Vande Voort JL, Croarkin PE. A Dose-Finding, Biomarker Validation, and Effectiveness Study of Transcranial Magnetic Stimulation for Adolescents With Depression. J Am Acad Child Adolesc Psychiatry. 2024 Sep 6:S0890-8567(24)01839–2. doi: 10.1016/j.jaac.2024.08.487. Epub ahead of print. PMID: 39245178; PMCID: PMC11882936.

Kappenman ES & Luck SJ. Best Practices for Event-Related Potential Research in Clinical Populations. Biol Psychiatry Cogn Neurosci Neuroimaging. 2016 1(2):110–115. PMID: 27004261; PMCID: PMC4797328.

Donchin E. Presidential address, 1980. Surprise!…Surprise? Psychophysiology. 1981 Sep;18(5):493–513. doi: 10.1111/j.1469-8986.1981.tb01815.x. PMID: 7280146.

Polich J. Updating P300: an integrative theory of P3a and P3b. Clin Neurophysiol. 2007 Oct;118(10):2128–48. doi: 10.1016/j.clinph.2007.04.019. Epub 2007 Jun 18. PMID: 17573239; PMCID: PMC2715154.

Owen AM, McMillan KM, Laird AR, Bullmore E. N-back working memory paradigm: a meta-analysis of normative functional neuroimaging studies. Hum Brain Mapp. 2005 May;25(1):46–59. doi: 10.1002/hbm.20131. PMID: 15846822; PMCID: PMC6871745.

Fitzgerald PB, Chen L, Richardson K, Daskalakis ZJ, Hoy KE. A pilot investigation of an intensive theta burst stimulation protocol for patients with treatment resistant depression. Brain Stimul. 2020 13(1):137–144. PMID: 31477542.

Beam W, Borckardt JJ, Reeves ST, George MS. An efficient and accurate new method for locating the F3 position for prefrontal TMS applications. Brain Stimul. 2009 Jan;2(1):50–4. doi: 10.1016/j.brs.2008.09.006. PMID: 20539835; PMCID: PMC2882797.

Caulfield KA, Fleischmann HH, Cox CE, Wolf JP, George MS, McTeague LM. Neuronavigation maximizes accuracy and precision in TMS positioning: Evidence from 11,230 distance, angle, and electric field modeling measurements. Brain Stimul. 2022 Sep-Oct;15(5):1192–1205. doi: 10.1016/j.brs.2022.08.013. Epub 2022 Aug 27. PMID: 36031059; PMCID: PMC10026380.

Sheehan DV, Lecrubier Y, Sheehan KH, Amorim P, Janavs J, Weiller E, Hergueta T, Baker R, Dunbar GC. The Mini-International Neuropsychiatric Interview (M.I.N.I.): the development and validation of a structured diagnostic psychiatric interview for DSM-IV and ICD-10. J Clin Psychiatry. 1998;59 Suppl 20:22–33;quiz 34-57. PMID: 9881538.

Zelazo PD, Anderson JE, Richler J, Wallner-Allen K, Beaumont JL, Conway KP, Gershon R, Weintraub S. NIH Toolbox Cognition Battery (CB): validation of executive function measures in adults. J Int Neuropsychol Soc. 2014 Jul;20(6):620–9. doi: 10.1017/S1355617714000472. Epub 2014 Jun 24. PMID: 24960301; PMCID: PMC4601803.

Kane RL, Short P, Sipes W, Flynn CF. Development and validation of the spaceflight cognitive assessment tool for windows (WinSCAT). Aviat Space Environ Med. 2005 Jun;76(6 Suppl):B183–91. PMID: 15943211.

Moore TM, Basner M, Nasrini J, Hermosillo E, Kabadi S, Roalf DR, McGuire S, Ecker AJ, Ruparel K, Port AM, Jackson CT, Dinges DF, Gur RC. Validation of the Cognition Test Battery for Spaceflight in a Sample of Highly Educated Adults. Aerosp Med Hum Perform. 2017 Oct 1;88(10):937–946. doi: 10.3357/AMHP.4801.2017. PMID: 28923143.

Gevins A, Cutillo B. Spatiotemporal dynamics of component processes in human working memory. Electroencephalogr Clin Neurophysiol. 1993 Sep;87(3):128–43. doi: 10.1016/0013-4694(93)90119-g. PMID: 7691540.

Keil A, Debener S, Gratton G, Junghöfer M, Kappenman ES, Luck SJ, …, Yee CM. Committee report: Publication guidelines and recommendations for studies using electro-encephalography and magnetoencephalography. Psychophysiology. 2014 Jan;51(1):1–21.

Gratton G, Coles MG, Donchin E. A new method for off-line removal of ocular artifact. Electroencephalogr Clin Neurophysiol. 1983 Apr;55(4):468–84.

Junghöfer M, Elbert T, Tucker DM, Rockstroh B. Statistical control of artifacts in dense array EEG/MEG studies. Psychophysiology. 2000 Jul;37(4):523–32.

Thigpen NN, Kappenman ES, Keil A. Assessing the internal consistency of the event-related potential: An example analysis. Psychophysiology. 2017 Jan;54(1):123–38.

Carrasco CD, Simmons AM, Kiat JE, Luck SJ. Enhanced Working Memory Representations for Rare Events. Psychophysiology. 2025 Mar;62(3):e70038. doi: 10.1111/psyp.70038. PMID: 40119595.

Cuthbert BN, Schupp HT, Bradley M, McManis M, Lang PJ. Probing affective pictures: attended startle and tone probes. Psychophysiology. 1998 May;35(3):344–7. doi: 10.1017/s0048577298970536. PMID: 9564755.

Jennings JR. Editorial policy on analyses of variance with repeated measures. Psychophysiology. 1987;24:474–5.

Keselman HJ. Testing treatment effects in repeated measures designs: An update for psychophysiological researchers. Psychophysiology. 1998;35(4):470–8.

Ebrahimzadeh E, Sadjadi SM, Asgarinejad M, Dehghani A, Rajabion L, Soltanian-Zadeh H. Neuroenhancement by repetitive transcranial magnetic stimulation (rTMS) on DLPFC in healthy adults. Cogn Neurodyn. 2025 Dec;19(1):34. doi: 10.1007/s11571-024-10195-w. Epub 2025 Jan 24. PMID: 39866659; PMCID: PMC11759757.

Heinze HJ, Luck SJ, Mangun GR, Hillyard SA. Visual event-related potentials index focused attention within bilateral stimulus arrays. I. Evidence for early selection. Electroencephalogr Clin Neurophysiol. 1990 Jun;75(6):511–27. doi: 10.1016/0013-4694(90)90138-a. PMID: 1693896.

Luck SJ, Kappenman ES, Fuller RL, Robinson B, Summerfelt A, Gold JM. Impaired response selection in schizophrenia: evidence from the P3 wave and the lateralized readiness potential. Psychophysiology. 2009 Jul;46(4):776–86. doi: 10.1111/j.1469-8986.2009.00817.x. Epub 2009 Apr 6. PMID: 19386044; PMCID: PMC2706937.

